# Pharmacological Ascorbate Induces Transient Hypoxia Sensitizing Pancreatic Ductal Adenocarcinoma to a Hypoxia Activated Prodrug

**DOI:** 10.1101/2024.05.13.593896

**Authors:** Shun Kishimoto, Daniel R. Crooks, Otowa Yasunori, Yamashita Kota, Kazutoshi Yamamoto, W. Marston Linehan, Mark Levine, Murali C Krishna, Jeffrey R Brender

## Abstract

Hypoxic tumor microenvironments pose a significant challenge in cancer treatment. Hypoxia-activated prodrugs like evofosfamide aim to specifically target and eliminate these resistant cells. However, their effectiveness is often limited by reoxygenation after cell death. We hypothesized that ascorbate’s pro-oxidant properties could be harnessed to induce transient hypoxia, enhancing the efficacy of evofosfamide by overcoming reoxygenation.

To test this hypothesis, we investigated the sensitivity of MIA Paca-2 and A549 cancer cells to ascorbate in vitro and in vivo. Ascorbate induced a cytotoxic effect at 5 mM that could be alleviated by endogenous administration of catalase, suggesting a role for hydrogen peroxide in its cytotoxic mechanism. In vitro, Seahorse experiments indicated generation of hydrogen peroxide consumes oxygen, which is offset at later time points by a reduction in oxygen consumption due to hydrogen peroxide’s cytotoxic effect.

In vivo, photoacoustic imaging showed ascorbate treatment at sublethal levels triggered a complex, multi-phasic response in tumor oxygenation across both cell lines. Initially, ascorbate generated transient hypoxia within minutes through hydrogen peroxide production, via reactions that consume oxygen. This initial hypoxic phase peaked at around 150 seconds and then gradually subsided. However, at longer time scales (approximately 300 seconds) a vasodilation effect triggered by ascorbate resulted in increased blood flow and subsequent reoxygenation. Combining sublethal levels of ascorbate with evofosfamide significantly prolonged tumor doubling time in MIA Paca-2 and A549b xenografts compared to either treatment alone. This improvement, however, was only observed in a subpopulation of tumors, highlighting the complexity of the oxygenation response.

---

One of the proposed biological functions of ascorbic acid (vitamin C) is to protect cellular components against free radicals generated during both normal metabolism and during periods of oxidative stress, either by direct reaction with aqueous free radicals[1, 2] or by recycling oxidized α-tocopherol (vitamin E).[3] However, the *in vivo* antioxidant activity of ascorbate is debated,[4] with some evidence suggesting it may act as a pro-oxidant in the presence of transition metals, potentially promoting oxidative stress and exerting adverse effects on biological systems.[5, 6]

The mechanism has been extensively investigated by Chen.et.al..[7, 8] and others.[9-11] In their proposed mechanism (Fig. 1),[12] the high concentration of ascorbate loses one electron to form Asc.-., which reduces a protein-centered metal and the reduced metal donates the electron to oxygen, forming active oxygen including superoxide with subsequent dismutation to H_2_O_2_. Several studies have reported the attenuation of treatment effects by catalase, confirming the pro-oxidant potential of ascorbate under these conditions. [9, 13] Consequently, under certain circumstances, ascorbate, acting as a reducing agent, can generate oxidants. Due to the low concentrations of free metals and the relatively low accessibility of metals in metalloproteins, this reaction is believed to occur only to an appreciable amount *in vivo* when concentrations in the millimolar range are achieved in the extracellular space through the intravenous administration of high-dose ascorbate, referred to as pharmacologic ascorbate. Oral delivery of ascorbate, even at high doses, is tightly controlled by the gastrointestinal system, limiting its bioavailability and preventing the attainment of these pro-oxidant concentrations.[14-16]

**Figure 1.**
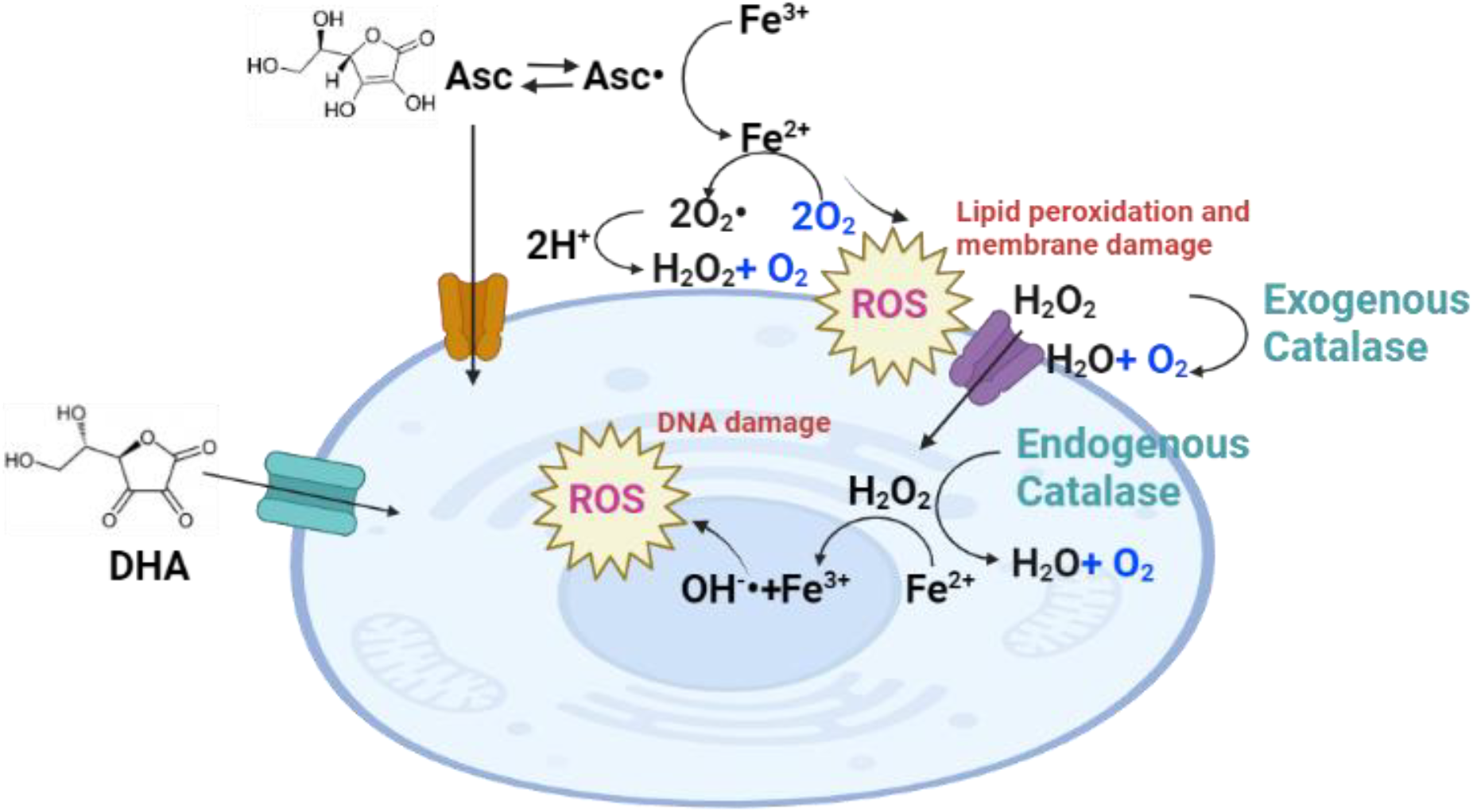
Simplified mechanism underlying the pro-oxidant cytotoxicity of pharmacological ascorbate in cancer cells focusing on oxygen production. At pharmacological concentrations, ascorbate donates an electron to reduce Fe3+/Cu2+ in the extracellular space to form Fe2+/Cu+. These reduced metals donate electrons to oxygen, producing hydrogen peroxide (H_2_O_2_) via superoxide while consuming oxygen. The hydrogen peroxide produced may be consumed during the Haber-Weiss reaction radical (OH•) (see reference [55] for other possible reactions). The net loss for the production of hydrogen peroxide is one molecule of oxygen. Intacellular hydrogen peroxide may then be decomposed by dismutation to generate one molecule of oxygen. Alternatively, two molecules of the ascorbate radical can combine to form dehydroascorbic acid (DHA). DHA is transported into cells via the glucose transporter GLUT1 and can be reduced back to ascorbate by a glutathione-dependent reaction catalyzed by dehydroascorbic acid reductase (DHAR) or by glutathione itself.

This intriguing duality in the behavior of ascorbic acid has sparked considerable interest among researchers in recent years, prompting a reevaluation of its physiological and pathological implications.[5] Although some aspects of the mechanism are incomplete,[17, 18] cancer cells show a unique vulnerability to the pro-oxidant effects of ascorbate.[8] Pharmacological ascorbate has been shown to exhibit selective pro-oxidant activity in tumor cells, leading to hydrogen peroxide-dependent cytotoxicity without adversely affecting normal cells. [7] The production of hydrogen peroxide by ascorbate-mediated reduction of iron and copper ions is a critical factor in this selective cytotoxicity. Tumor cells have higher levels of labile iron[19] and lower catalase activity[11] compared to normal tissue, making them more susceptible to the pro-oxidant effects of ascorbate Furthermore, the high metabolic rate in tumor cells also contributes to endogenously higher oxidative stress levels, further enhancing their vulnerability. [20]

The unique vulnerability of cancer cells to the pro-oxidant effects of ascorbate has been leveraged in combination with other anti-cancer treatments. Preclinical studies have demonstrated promising results,[10, 11, 21-24] such as the combination of ascorbate with gemcitabine resulting in greater decreases in tumor volume and weight compared to gemcitabine alone in a mouse model of pancreatic cancer.[22] Similarly, multiple preclinical investigations have shown the potential benefit of combining ascorbate with radiation therapy.[13, 25, 26] Early phase clinical trials have evaluated the safety and tolerability of combining pharmacological ascorbate with standard therapies like radiation and chemotherapy.[23, 27-31] Some of these studies have reported improved outcomes,[26, 29, 32, 33] but definitive efficacy remains to be established in prospective randomized trials.[32] The enhanced efficacy is attributed to ascorbate’s ability to increase oxidative stress and potentiate the cytotoxic effects of these therapies in the tumor microenvironment.

While oxidative stress caused by ascorbate continues to be investigated for its cytotoxic effects, another aspect of ascorbate pro-oxidant pharmacology can potentially be utilized for treatment. The oxidation of ascorbate initially consumes oxygen,[7] which, under the poorly developed vasculature of the tumor microenvironment, can lead to localized oxygen depletion. By using sublethal doses of ascorbate to induce transient hypoxia without direct cytotoxicity, it is possible to isolate the synergistic effect of ascorbate and evofosfamide from the toxicity of ascorbate alone. This approach allows us to exploit the oxygen-consuming properties of ascorbate to create a unique opportunity for the advancement of hypoxia-targeting treatments. These drugs aim to improve the effectiveness of therapies like radiation and chemotherapy by exerting their cytotoxic effects specifically in hypoxic areas of the tumor where the effectiveness of these agents is reduced.[34, 35] We show that by exploiting the oxygen-consuming properties of ascorbate, the effectiveness of the hypoxia targeting drug evofosamide can.be enhanced to target tumors that are only mildly sensitive to evofosfamide alone. Critically, the effect is transient, and the vasodilation effect induced by ascorbate at longer intervals reoxygenates the sample once evofosfamide has taken effect, providing an opportunity for further therapeutic intervention in an oxygenated environment.

## Materials and methods

### *In vivo* metabolic flux assay

The oxygen consumption rate (OCR) and extracellular acidification rate (ECAR) were analyzed using an XF96 Extracellular Flux Analyzer (Agilent Technology). Twenty thousand cells were plated into each well of a 96-well plate and cultured overnight. During the measurement of OCR and ECAR, cells were treated with either ascorbate (2.5, 5, 10, and 20 mM) or hydrogen peroxide (25, 50, 100, and 200 µM) at the 30-minute time point and with catalase (100 µg/ml) at the 60-minute time point. OCR was calculated based on a three compartment model to account for the effects of oxygen diffusion during the experiment, which led to a cycling pattern reflecting the transient changes in oxygen availability and consumption over time.[36] (Supplementary Figure 2).

### Western blotting

MIA PaCa-2 and A549 tumor tissues were excised and homogenized in T-PER Tissue Protein Extraction Reagent (Thermo Fisher Scientific). Protein concentrations were measured using the bicinchoninic acid assay (BCA protein assay, Thermo Fisher Scientific). Glut-1 and catalase proteins were separated on a 4% to 20% Tris-Glycine gel (Life Technologies) by SDS-PAGE and transferred to a nitrocellulose membrane. The membranes were blocked for 1 hour in blocking buffer (3% nonfat dry milk in 0.1% Tween 20/TBS), which was then replaced by the primary antibody (1:500–1:1,000), diluted in blocking buffer, and incubated for 1 hour at room temperature. The membranes were then washed three times in washing buffer (0.1% Tween 20/TBS). The primary antibody was detected using the appropriate horseradish peroxidase-conjugated secondary antibody and measured by the Fluor Chem HD2 chemiluminescent imaging system (Alpha Innotech Corp.). Density values for each protein were normalized to HSC70.

### Animal experiments

All animal experiments were conducted in compliance with the Guide for the Care and Use of Laboratory Animal Resources (National Research Council, 1996), and the experimental protocols were approved by the National Cancer Institute Animal Care and Use Committee (RBB-159-2SA). MIA PaCa-2 and A549 cells were routinely cultured in RPMI 1640 with 10% fetal calf serum. Tumors were formed by injecting 3 × 106 cells subcutaneously into the right hind legs of female athymic mice. Tumor-bearing mice were treated with intraperitoneal injections of ascorbate (2 g/kg) every other day for 7 days when the tumor size reached approximately 40 mm^3^. For combination treatment experiments, ascorbate (2 g/kg) was injected one hour after the intraperitoneal injection of evofosfamide (50 mg/kg) three times weekly to ensure intratumor distribution of evofosfamide.

### Cell Viability assay

MIA PaCa-2 and A549 cells were seeded into 96-well plates at a density of 10,000 cells/well 24 h before the addition of ascorbate and/or catalase. After the drug addition, the cells were incubated for 1 hour at 37°C in a standard CO2 tissue culture incubator. Cell cytotoxicity was assessed using the CellTiter-Glo Luminescent Cell Viability assay (Promega Corporation, Madison, USA). The luminescent signal was measured using the GloMax luminescence detector. Data were presented as proportional viability (%) by comparing the treated group with the untreated cells, the viability of which is assumed to be 100%.

### Photoacoustic imaging experiments

Tumors were scanned with the Visual Sonics Vevo®LAZR System (FUJIFILM VisualSonics Inc., Canada) using a 21-MHz linear array transducer system (central frequency) integrated with a tunable nanosecond pulsed laser. The system was calibrated by the manufacturer before use. During the scan, mice were anesthetized using isoflurane (1.5–2.5%). Respiration was continuously monitored to ensure data reproducibility and animal well-being. The tumor area in the sagittal plane of the leg was manually determined from concurrently acquired ultrasound images. In this study, the experiments were performed on a fixed plane depicting the center of the tumor to achieve better temporal resolution. For O2 status assessments, photoacoustic images in the tumor area were collected with OxyHemo Mode (wavelength 750 nm / 850 nm) for 10 min. Oxygen saturation of hemoglobin (sO2) and total hemoglobin values in the tumor area were calculated using the OxyZated™ tool based on a previously reported and tested algorithm. [37] In order to reduce the noise level, the threshold of total hemoglobin in relation to the maximum possible level was set to 20% for all experiments. Values below the threshold were discarded and appear black in all images.

### Statistical analyses

Data were expressed as the means ± standard error. The significance of the differences between groups was analyzed using the Student t-test. p < 0.05 was considered statistically significant.

## Results

### MIA Paca-2, but not A549, tumors are sensitive to ascorbate *in vitro* and *in vivo*

To further investigate the pro-oxidant activity of pharmacological ascorbate, we conducted experiments using two cell lines: a cell line whose xenografts are known to be sensitive to pharmacological ascorbate (MIA Paca-2)[11] and a lung carcinoma cell line suspected to be resistant based on *in vitro* cell viability assays.[38] We first assessed their cytotoxic responses to ascorbate in vitro using an intracellular ATP viability assay, confirming MIA Paca-2’s higher sensitivity under the same conditions (Fig. 2A). To confirm the results *in vivo*, we examined ascorbate’s effects on tumor size progression in xenografts of the two cell lines (Fig. 2B). Mice received intraperitoneal injections of 2 g/kg ascorbate every other day for 7 days, starting when tumors reached a size of 40 mm^3^ due to the expected weak cytotoxic effect of ascorbate. Consistent with the *in vitro* results, treatment resulted in modest anti-tumor activity in MIA Paca-2 tumors but had no observable effect on A549 tumors.

**Figure 2:**
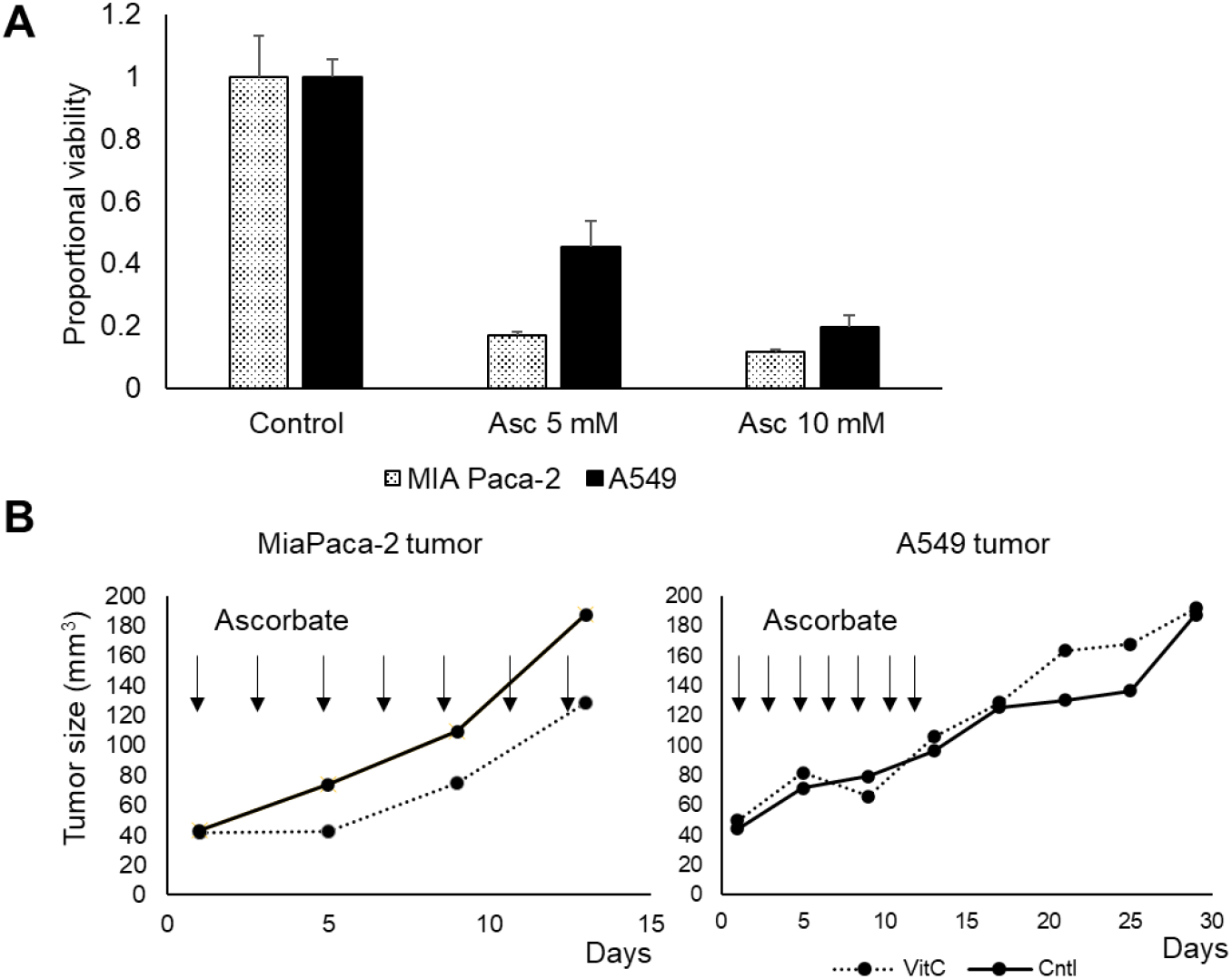
Ascorbate decreases in vitro viability and moderately suppresses tumor growth in MiaPaCa-2 but not in A549 tumors. **(A)** Viability in MiaPaCa-2) and ascorbate resistant cells (A549) upon ascorbate treatment based on a ATP production assay **(B)** Tumor growth in MiaPaCa-2 and A549 tumors a treatment with intraperitoneal injections of ascorbate (2 g/kg) every other day for 14 days after the tumor size reached approximately 40 mm^3^

### Differential Catalase Activity in MIA Paca-2 and A549 Mediate Ascorbate Sensitivity

The sensitivity of cancer cells to pharmacological ascorbate treatment may be influenced by their ability to reduce hydrogen peroxide or uptake dehydroascorbic acid (DHA), [10, 26] the oxidized form of ascorbate primarily imported into cells via the GLUT1 transporter (Fig.1). [10, 39] To investigate the factors contributing to the differential sensitivity between MIA Paca-2 and A549 cells, we performed Western blotting to assess the levels of catalase, an enzyme that decomposes hydrogen peroxide, and glucose transporter-1 (GLUT1). The results showed that while catalase levels were lower in ascorbate-sensitive MIA Paca-2 cells, GLUT1 levels were similar between both cell lines (Fig. 3A). This suggests that the sensitivity to ascorbate treatment was at least partially attributed to differences in catalase activity. To further support this hypothesis, we demonstrated that the addition of 100 μg/ml exogenous catalase eliminated the cytotoxic effect of 5 mM ascorbate in both cell lines( Fig. 3B). These findings indicate that the treatment effect of pharmacological ascorbate observed in vivo is mainly caused by extracellular hydrogen peroxide and is largely independent of the intracellular import of ascorbate or DHA. [10]

**Figure 3:**
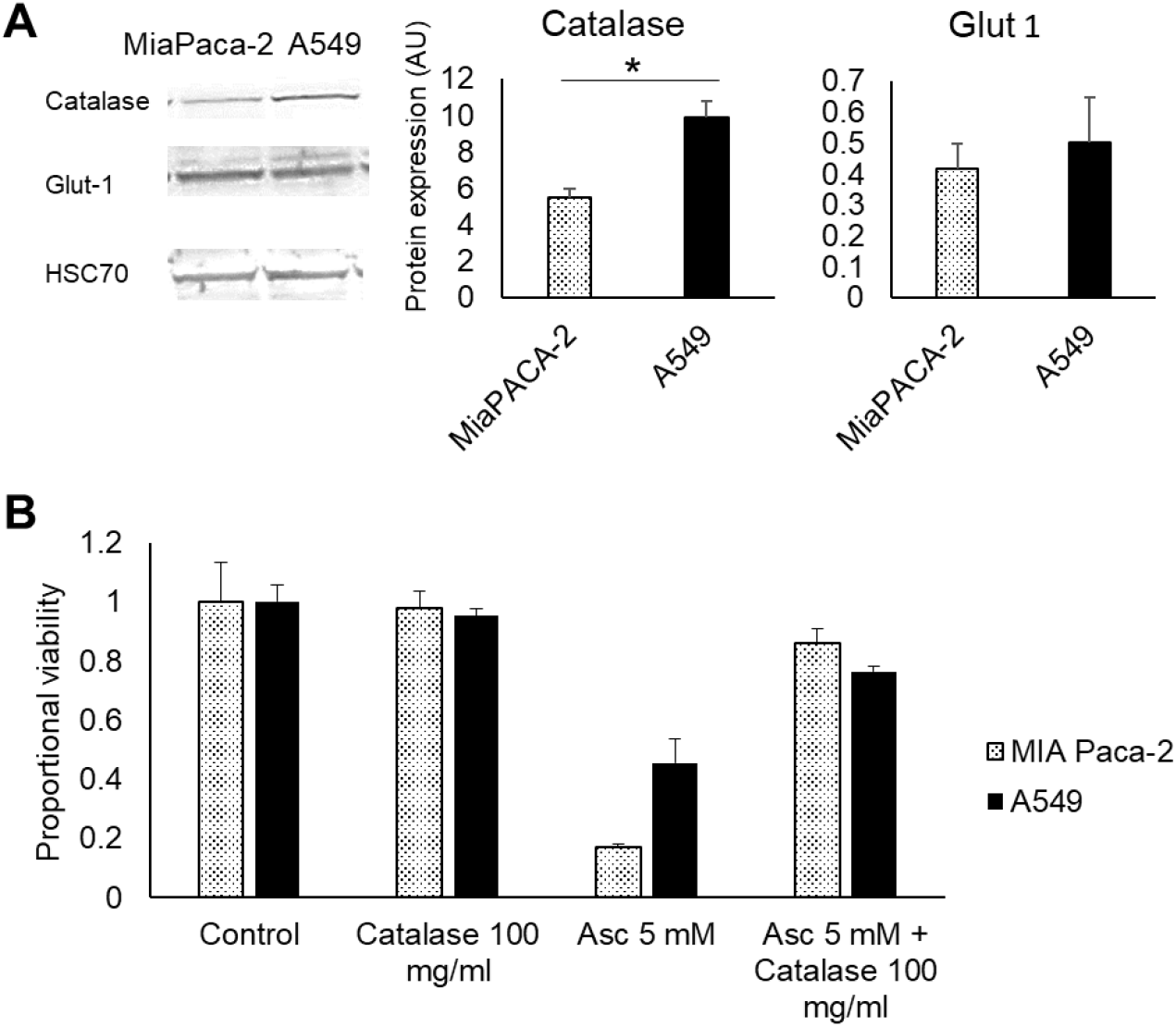
In vitro toxicity is suppressed by catalase. **(A)** Western blot of catalase, GLUT1, and the HSC70 marker **(B)** Viability after addition of 100 µg/ml exogenous catalase

### Hydrogen Peroxide Increases Oxygen Consumption at Short Timescales and Decreases Oxygen Consumption at Longer Timescales

To test this hypothesis, we evaluated the production of hydrogen peroxide indirectly by assessing the oxygen consumption rate (OCR) after ascorbate treatment using a metabolic flux analyzer *in vitro*. Hydrogen peroxide is produced by a two-step process in which oxygen is initially consumed during the Fenton reaction to produce hydrogen peroxide. The hydrogen peroxide produced by the Fenton reaction is then consumed during the Haber-Weiss cycle to generate oxygen and the highly reactive hydroxyl radical.

The net loss for both reactions is one molecule of oxygen.(Fig. 1). In line with this scheme, ascorbate treatment as measured by metabolic flux analysis initially induced a pronounced increase in oxygen consumption rate (OCR) in both MIA Paca-2 and A549 cells (Fig. 4A), likely reflecting the predominance of the oxygen consuming Fenton reaction step at early timepoints. However, this boost was followed by a rapid decline in OCR, which can be attributed to the emergence of the oxygen producing Haber-Weiss cycle. Subsequent catalase treatment to remove hydrogen peroxide and stop the Haber-Weiss cycle reversed the decline, and even elevated OCR above baseline (Fig. 4A).

**Figure 4:**
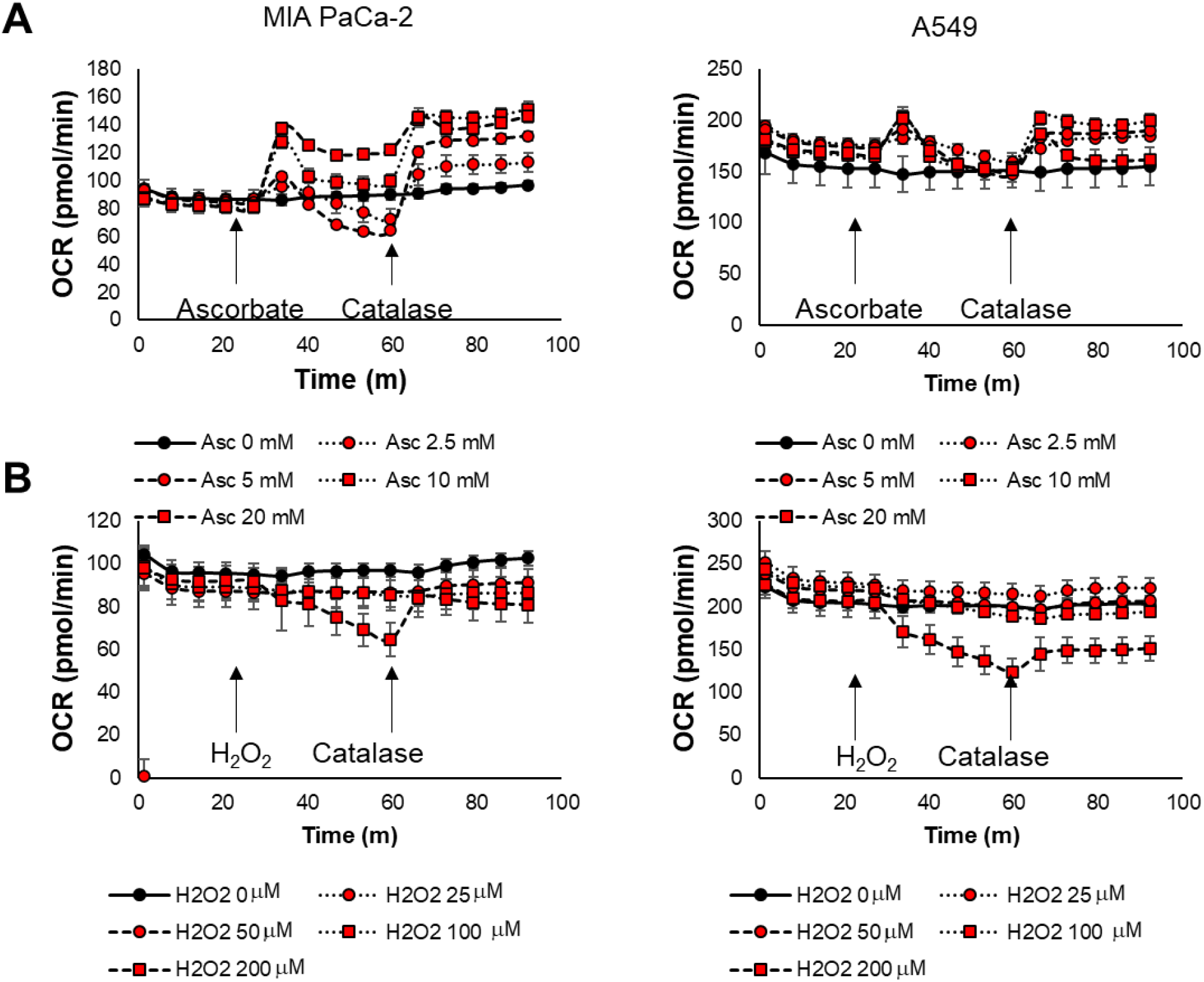
Oxygen consumption rate (OCR) calculated from a three compartment model after the addition of ascorbate followed by catalase (A) or hydrogen peroxide followed by catalase using a Glycolytic Rate assay kit on a Seahorse XF96e analyzer.

These results suggest that pharmacological ascorbate causes hydrogen peroxide production, resulting in an upregulated consumption of oxygen regardless of the cell type. Ascorbate treatment also reduced the rate of extracellular acidification (ECAR), indicating suppressed glycolysis (Fig. 4A). Catalase addition attenuated this decrease, suggesting that hydrogen peroxide hinders both glycolysis and anaerobic fermentation. However, unlike the enhancement of the OCR, this effect persisted even after catalase treatment (Supplementary Figure 2).

To confirm the role of hydrogen peroxide in damaging both mitochondrial respiration and anaerobic fermentation, we performed the same set of experiments using hydrogen peroxide instead of ascorbate (Fig. 4B) As anticipated, the initial OCR increase seen with ascorbate was absent due to the direct provision of hydrogen peroxide, which bypasses the oxygen-consuming Fenton reaction. Although the initial OCR response differed, the subsequent changes in both OCR and ECAR closely mirrored those observed with ascorbate and catalase treatment (Fig. 4B and Supplementary Figure 2). After hydrogen peroxide injection, OCR initially decreased but was quickly restored by catalase. Similarly, ECAR decreased and remained down even after catalase treatment, reflecting a persistent effect on glycolysis possibly due to the depletion of NAD+.[7, 40] These experiments demonstrated that pharmacological ascorbate increases oxygen consumption due to enhanced production of hydrogen peroxide. Notably, the baseline OCR was higher in A549 cells, and the extent of the OCR increase after ascorbate treatment was relatively higher in MIA PaCa-2 cells. This result aligns with the lower catalase activity and higher ascorbate sensitivity of MIA PaCa-2 cells.

### Ascorbate Generates Transient Hypoxia In Vivo at Short Time Scales and Increases Blood Flow at Longer Timescales

To validate the *in vivo* impact of ascorbate on tumor oxygenation, we employed photoacoustic imaging (PAI) in both MIA Paca-2 and A549 tumors (Fig. 5). PAI offers a unique advantage by providing real-time visualization and quantification of both oxyhemoglobin and total hemoglobin within living tissues. Briefly, through spectral analysis of hemoglobin at specific wavelengths (750 nm and 850 nm), PAI allows the non-invasive tracking of changes in blood oxygenation and blood volume dynamics within tumors.[41] As predicted oxygen saturation (sO2) decreases in both tumor types following ascorbate injection indicating a transient period of hypoxia (Fig. 5B). This transient hypoxia, lasting 5-6 minutes after ascorbate injection, is evidence that ascorbate does indeed trigger enhanced oxygen consumption within tumors *in vivo*, mirroring the effects observed in our *in vitro* experiments (Fig. 4B). Following the hypoxic phase, a substantial increase in total hemoglobin levels was observed, exceeding baseline levels for approximately 5 minutes (Fig. 5B). This biphasic response, characterized by initial hypoxia followed by vasodynamic changes that increase blood flow, was absent in MIA Paca-2 tumors treated with vehicle alone. Independent of its metabolic effects, pharmacologic ascorbate possesses vasodilatory properties.[42-44] This vasodilation could lead to an influx of blood into the tumor following the initial hypoxia, consistent with the observed increase in total hemoglobin.

**Figure 5:**
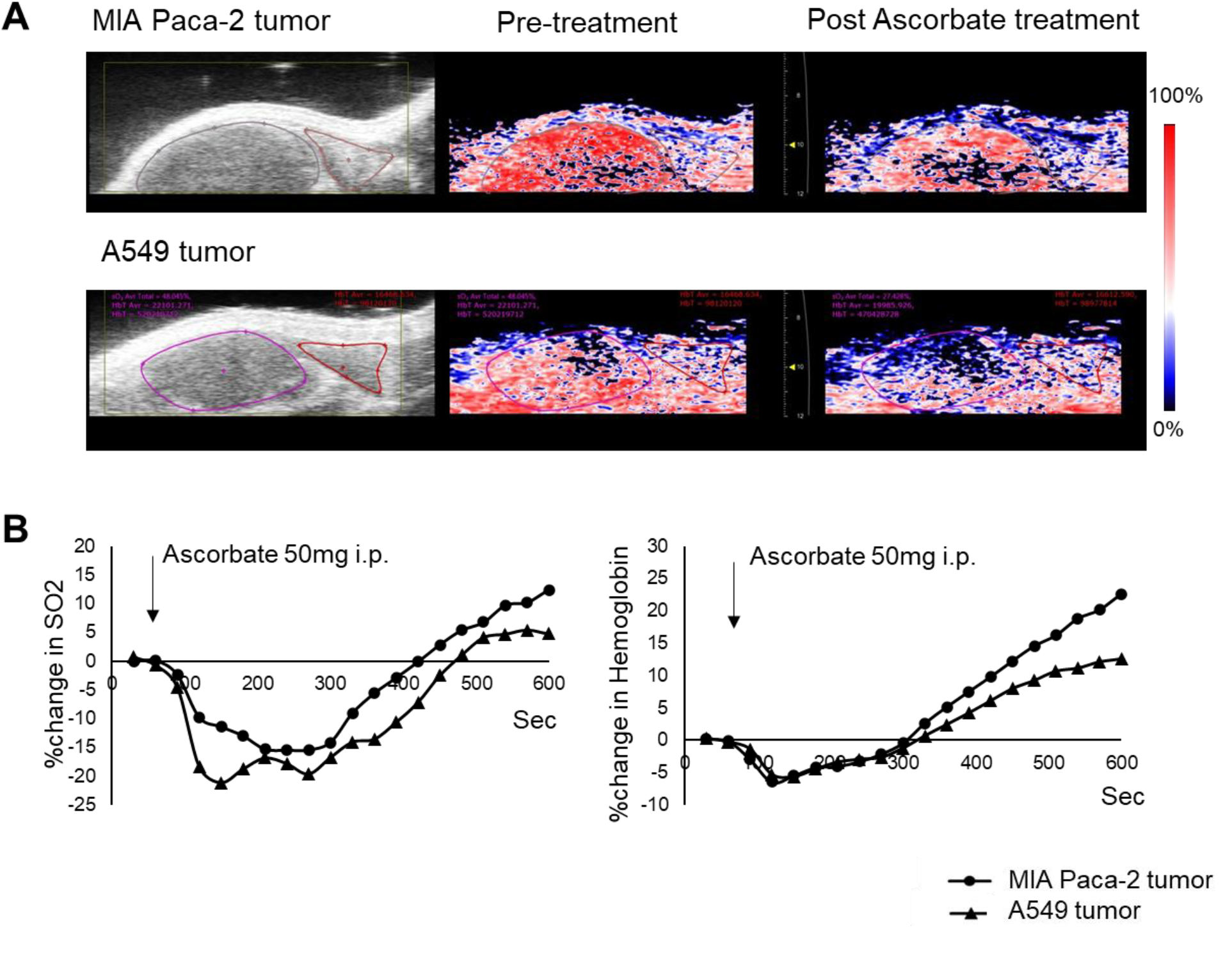
(A) Photoacoustic imaging of a MiaPaCa-2 and A549 tumor. (B) Percent change in oxygen saturation of hemoglobin (sO2) and total hemoglobin values for the regions of interest indicated in (A)

### Transient Hypoxia from Ascorbate Treatment Enhances the Efficacy of Evofosfamide

The induction of hypoxia by ascorbate, independent of its cytotoxic effect, might be useful to enhance the treatment efficacy of other hypoxia-dependent anti-cancer therapies, such as evofosfamide (Evo). [45] Evofosfamide is a rapid acting, hypoxia-activated prodrug that undergoes enzymatically activated fragmentation to release bromo-isophosphoramide mustard (Br-IPM) in the absence of oxygen.[25, 34] To investigate the synergistic effect of Evofosfamide and pharmacological ascorbate, we conducted tumor growth experiments using MIA Paca-2 and A549 tumor models. Four groups were examined: a control, an ascorbate treatment group (2 g/kg intraperitoneal injection three times per week), Evo treatment (50 mg/kg intraperitoneal injection three times per week), and a combination treatment group (Evo treatment one hour prior to ascorbate treatment).

In both MIA Paca-2 and A549 tumor models, neither evofosfamide nor ascorbate monotherapy had a significant impact on tumor growth, with no statistically significant difference in tumor doubling time (TDT) compared to the control group. This lack of response may be attributed to the later treatment start (when tumors reached approximately 800 mm^3^) compared to other studies, which was necessary to ensure accurate size measurements.[11] However, further analysis revealed that a small subpopulation of tumors in both cell lines responded favorably to the combination of evofosfamide and ascorbate. This subpopulation, characterized by a TDT more than twice the mean value, was three times larger in the combination treatment group compared to either monotherapy group. The presence of this responsive subpopulation resulted in a statistically significant difference in mean TDT between the combination therapy and control groups, despite the lack of response to either treatment alone. Notably, no systemic toxicity was observed in any of the treatment groups throughout the experiments. These findings suggest that the combination of ascorbate and evofosfamide can enhance treatment efficacy in a subset of tumors, potentially by leveraging ascorbate-induced transient hypoxia to activate evofosfamide in otherwise non-responsive tumors.

## Discussion

Hypoxic tumor microenvironments, characterized by oxygen deprivation, are a significant hurdle to successful cancer treatment.[34] These regions foster the development of chemotherapy-resistant[46] and immunosuppressive[47] cell populations, ultimately contributing to treatment failure and poor prognosis across various malignancies. Consequently, targeted therapies capable of addressing these hypoxic niches hold immense promise for improving patient outcomes.

Hypoxia-activated prodrugs have emerged as a novel strategy in this fight. These unique therapeutic agents remain inactive in well-oxygenated tumor areas but undergo a bioreductive conversion within hypoxic regions, transforming into potent cytotoxic alkylating agents.[25, 34, 35] This targeted mechanism offers a distinct advantage over conventional chemotherapies by specifically targeting hypoxia-driven chemoresistance and eliminating resistant cells residing in oxygen-depleted sanctuaries.

However, despite the theoretical elegance of this approach, clinical results for evofosfamide, the leading hypoxia activated prodrug candidate, have been mixed. The MAESTRO Phase III trial for unresectable pancreatic ductal adenocarcinoma, while demonstrating a statistically significant improvement in progression-free survival with evofosfamide/gemcitabine combination therapy compared to gemcitabine alone, failed to meet the primary endpoint of overall survival benefit.[48] This highlights the limitations of evofosfamide monotherapy and underscores the need to optimize its clinical application.[49]

A critical factor influencing its efficacy is the dynamic nature of the tumor microenvironment.[50] While evofosfamide effectively eliminates hypoxic cells, this very action can paradoxically lead to tumor reoxygenation.[51] Extensive cell death triggered by the drug significantly reduces cellular oxygen consumption within the tumor. However, in the absence of concomitant changes in oxygen supply, this decline in demand can elevate intra-tumoral oxygen levels, effectively reoxygenating previously hypoxic regions.[51] Consequently, reoxygenation can limit the activation and efficacy of evofosfamide in previously targeted areas.

Strategies to mitigate reoxygenation, through combination therapies or microenvironmental manipulation, are crucial for maximizing the therapeutic potential hypoxia active prodrugs. Transient hypoxia can be induced by either decreasing the oxygen supply or increasing the oxygen demand. The vasodilator hydralazine has been used to reduce tumor blood flow and improve evofosfamide efficacy through the “steal” phenomenon, in which atonal immature tumor vasculature fails to dilate in coordination with normal vasculature.[52] In this context, vasodilators have a limited therapeutic range and their effective use is dependent on the steal phenomenon and will not work in tumors with highly developed vasculature.[50] In preclinical models, transient hypoxia has also been induced with an intravenous bolus of pyruvate. While pyruvate improved evofosfamide treatment in several PDAC mouse models, [45, 53] induction of transient hypoxia by pyruvate relies on OXPHOS and its potential in highly glycolytic tumors is untested. Consequently, each of these methods have some limitations that may limit their translational potential. Additionally, induction of transient hypoxia in any form will create an immunosuppressive and radiation resistant environment with potentially deleterious consequences for combination therapy.

Our study shows short-term exposure to pharmacological doses of ascorbate can generate transient hypoxia via heightened oxygen consumption rates (OCR) *in vitro* (Fig. 4) and *in vivo* (Fig. 5), thereby sensitizing otherwise minimally responsive[50] tumors to hypoxia-activated prodrugs like evofosfamide (Fig. 6). Interestingly, enhanced oxygen consumption was evident in both ascorbate sensitive MIA Paca-2 and resistant A549 cells (Fig. 4) and tumors from both cell line were susceptible to combination therapy, indicating the transient hypoxia effect is largely independent of direct ascorbate cytotoxicity despite the known link between hydrogen peroxide production and cytotoxicity.[54]

**Figure 6:**
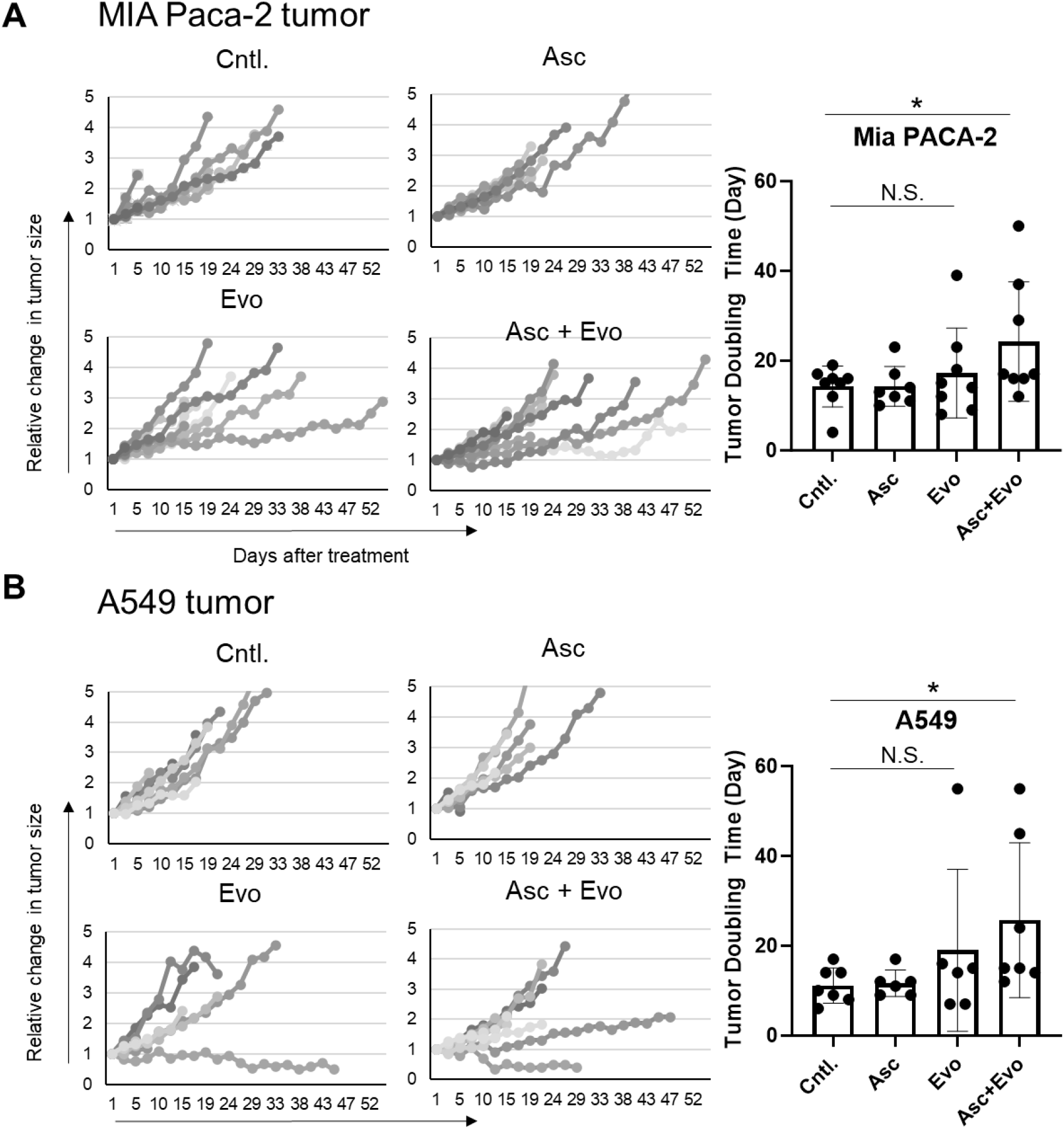
Left: Tumor growth in MiaPaCa-2 (A) and A549 (B) tumors after a 7 day regimen of treatment with intraperitoneal injections of ascorbate (2 g/kg) and/or evofosfamide (50 mg/kg) every other day. Right: Tumor doubling times calculated from the tumor growth curves

The complex oxygen response seen with photoacoustic imaging aligns with ascorbate’s dual capacity to stimulate reactive oxygen species (ROS) production and vasodilation. The initial hypoxia mirrors the heightened oxygen consumption and reduced tumor oxygenation levels demonstrated in the *in vitro* and *in vivo* experiments (Figs. 4 and 5). This transient hypoxia sensitizes tumors to hypoxia-targeted therapies like evofosfamide. However, the subsequent increase in total hemoglobin and oxygenation mediated by ascorbate’s vasoactive properties could enable reoxygenation and limit prolonged hypoxia. Importantly, the relative impacts of these competing effects likely vary across individual tumors. As noted, only a subpopulation of tumors responded favorably to evofosfamide alone or in combination with ascorbate, possibly due to poor penetration in the poorly vascularized core region of the tumor.[27, 50] To further enhance the therapeutic efficacy, the ascorbate dose could be increased to take advantage of direct ascorbate toxicity at higher doses and a larger hypoxic effect. Tailoring combination schedules to the timing and durations of hypoxia versus reoxygenation elicited by ascorbate will be vital to synchronize treatments and combat therapy resistance. Additional imaging-guided studies mapping intratumoral oxygen fluctuations on a patient-specific basis could empower more precise deployment of hypoxia-activating agents.[34]

---

**Supplementary Figure 1:**
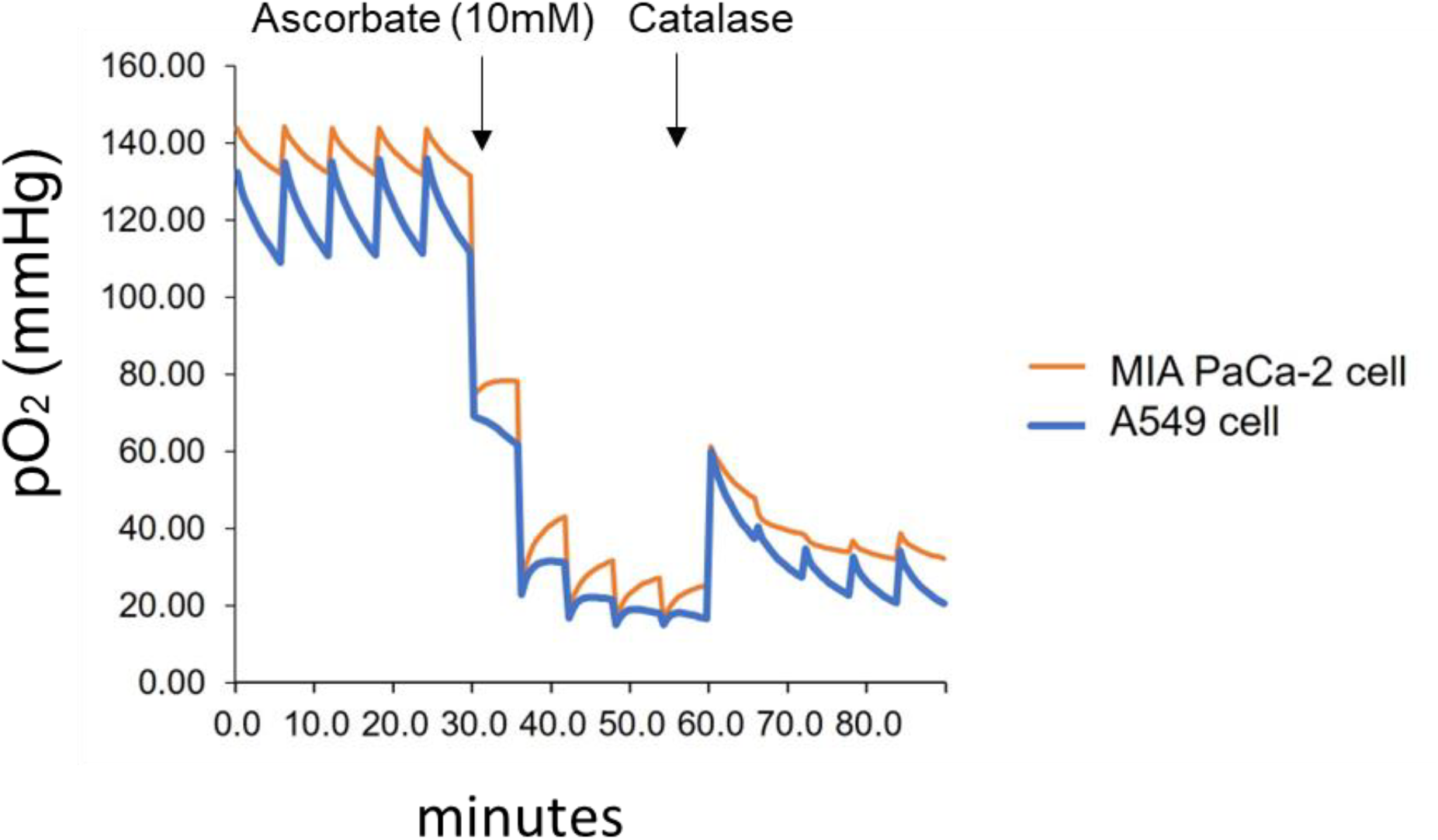
Representative oxygen profile uncorrected for diffusion

**Supplementary Figure 2:**
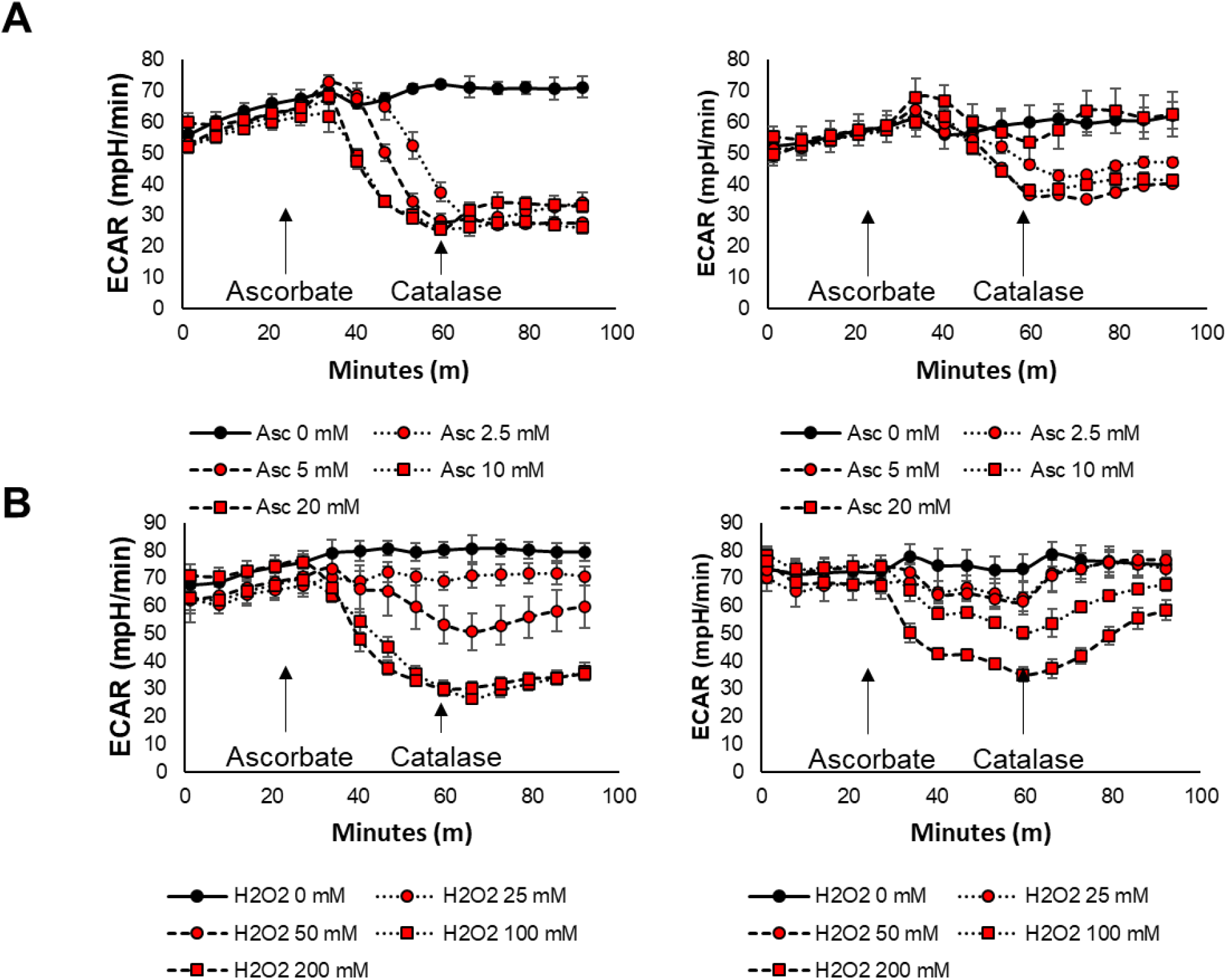
Extracellular Acidification Rates (Rates) for the OCR plotted in Figure 3. Add seconds to x axis or indicate x axis is in seconds.

